# Molecular and cellular architecture of the larval sensory organ in the cnidarian *Nematostella vectensis*

**DOI:** 10.1101/2021.05.10.443235

**Authors:** Callum Teeling, Eleanor Gilbert, Siffreya Pedersen, Nathan Chrismas, Vengamanaidu Modepalli

## Abstract

The apical pole of eumetazoan ciliated larvae acts as a neurosensory structure and is principally composed of sensory-secretory cells. Cnidarians like the sea anemone *Nematostella vectensis* are the only non-bilaterian group to evolve ciliated larvae with a neural integrated sensory organ that is likely homologous to bilaterians. Here, we uncovered the molecular signature of the larval sensory organ in *Nematostella* by generating a transcriptome of the apical tissue. We characterised the cellular identity of the apical domain by integrating larval single-cell data with the apical transcriptome and further validated this through in-situ hybridisation. We discovered that the apical domain comprises a minimum of 6 distinct cell types, including apical cells, neurons, peripheral flask-shaped gland/secretory cells, and undifferentiated cells. By profiling the spatial expression of neuronal genes, we showed that the apical region has a unique neuronal signature distinct from the rest of the body. By combining the planula cilia proteome with the apical transcriptome data, we revealed the sheer complexity of the non-motile apical tuft. Overall, we present comprehensive spatial/molecular data on the *Nematostella* larval sensory organ and open new directions for elucidating the functional role of the apical organ and larval nervous system.

## Introduction

The majority of marine benthic invertebrates during their early development progress through a planktonic life phase, a ciliated larva with an apical organ [1]. Several behavioural studies have demonstrated that ciliated larvae use the sensory organ at the apical pole to sense environmental cues and modulate their swimming behaviour [2-4]. The apical pole of the larvae is enriched with several flask-shaped cells, usually with an apical tuft of non-motile cilia **(Fig 1)** [1, 5-11]. Besides the flask-shaped apical cells, the sensory (photosensitive and mechanosensory) and secretory/gland cells are also scattered around the apical pole and are likely associated with the sensory-ciliomotor nervous system. For instance, in bilaterian trochophore larvae such as the mollusc *Ischnochiton hakodadensis* **(Fig 1F)** [12, 13] and the annelid *Malacoceros fuliginosus*, the apical pole possesses several sensory cells positive for serotonin and neuropeptide FMRFamide [14]. Similarly, the marine annelid *Platynereis dumerilii* is also equipped with photosensitive, mechanosensory and peptidergic cell types alongside an apical tuft [15-18].

**Figure 1:**
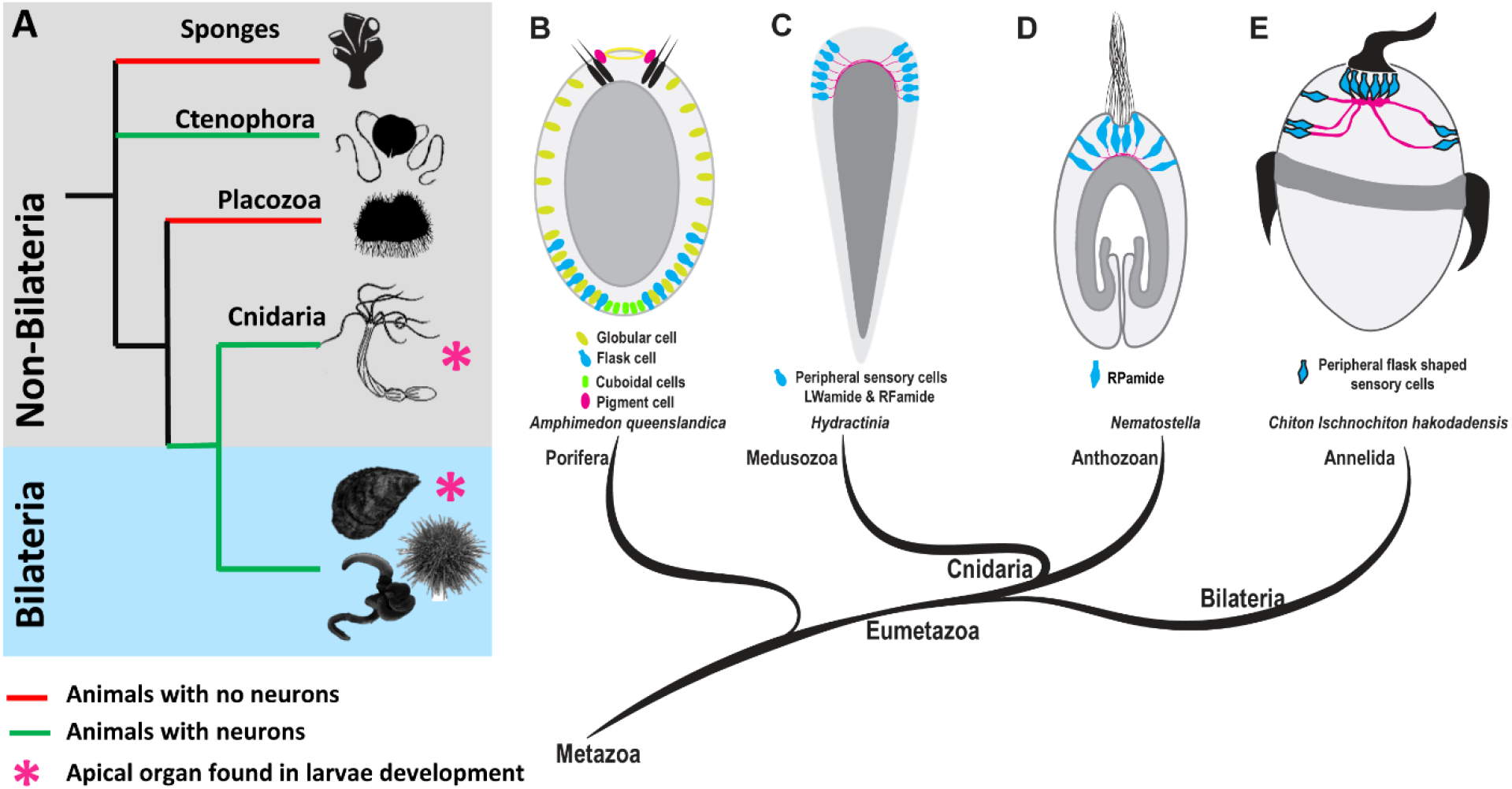
Origin and evolution of nervous system and apical organ. **(A)** A brief overview of evolutionary relationships in the animal kingdom. Porifera and Placozoa do not have defined neurons. Among non-bilaterians, Cnidaria and Ctenophora have well defined neurons. **(B)** Sponge larvae have ciliated photoreceptor cells capable of light sensing and other peripheral cell types such as flask and cuboidal cells. **(C & D)** Among non-bilaterians, a true larval apical organ with integrated neurons is only found in cnidarians. **(B-E)** Schematic drawings of surface-contacting flask-shaped cells in the ciliated larvae of different marine phyla. Schemes were drawn based on the following primary data: **(B)** Nakanishi, N., et al. 2015 [33], Richards, G. S. and B. M. Degnan 2012 [28], Ueda, N., et al. 2016 [34], **(C)** Gajewski, M., et al.1996 [24], Katsukura, Y., et al. 2004 [26], **(D)** Zang, H. and N. Nakanishi 2020 [22], **(E)** Voronezhskaya, E. E., et al. 2002 [12], Nezlin, L. P. and E. E. Voronezhskaya 2017 [13].

Among non-bilaterians, a larval sensory organ with integrated neurons is only found in cnidarians **(Fig 1)** [19-21]. In anthozoans, like the sea anemone *Nematostella*, the apical pole displays several flask-shaped apical cells with a ciliated tuft and RPamide-positive sensory cells **(Fig 1E)** [22]. In hydrozoans, the apical pole is highly enriched with LWamide and RFamide neuropeptide expressing cells (**Fig 1D**) [23-26]. However, unlike anthozoans, the hydrozoans lack an apical organ-like sensory structure with a ciliated tuft [47]. Strikingly, animals devoid of neurons, such as the sponge *Amphimedon queenslandica*, also bear a set of sensory and secretory cells (globular/mucous cells) in the anterior region of the larvae [33, 56], which are likely involved in modulating larval behaviour **(Fig 1 B)** [27-30]. However, the advent of neuronal coordination of motile cilia is strategic in increasing the efficiency of sensory-to-motor transformation [31]. If ciliated larvae were fundamental to the origin of the first eumetazoans [32], the origin of ciliated larvae with a neural integrated sensory organ is likely allied to the evolution of the first nervous system in eumetazoans (**Fig 1 A-D)**.

The morphology of the apical organ in cnidarian larvae is comparable to those of bilaterian larvae, indicating that the common ancestor of these two groups progressed through a free-swimming larval stage with a true larval apical organ and associated neurons [35, 36]. Earlier studies by Sinigaglia et al. have identified a range of apical organ genes by blocking FGF signalling during early development [2]. Comparative gene expression studies between the cnidarian *Nematostella* and bilaterian ciliated larvae revealed a strong resemblance in molecular topography around the apical pole [15, 37-39], suggesting that the apical organ is likely an evolutionarily conserved larval structure. If so, to what extent is the larval sensory organ functionally homologous among cnidarians and bilaterians? This demands functional evidence from cnidarians, which in turn requires in-depth molecular profiling of the cnidarian apical domain and its cellular composition.

Cnidarians hold a key phylogenetic position for understanding nervous system evolution and their larval sensory-ciliomotor nervous system provides a window to look into the primordial neurotransmission system. To advance this, we aimed to map the apical enriched cell types and their gene expression profiles in the anthozoan cnidarian *Nematostella vectensis. Nematostella* is a well-established molecular model species that has been at the centre of fundamental discoveries in the development and evolution of the nervous system in non-bilaterian metazoans, making it a suitable model for the current study [19-21]. Here, we reveal the expression profile of the larval sensory organ by performing transcriptomics on apical tissue that was separated from the rest of the larval body **(Fig 2A)**. Further, by integrating our tissue-specific (apical/body) transcriptome data with *Nematostella* larval single-cell RNA-seq data [40], we identified the larval cell types enriched in the body and apical regions and their gene expression profiles. Next, to gain insights into the non-motile apical tuft, we integrated the cilia proteome data for *Nematostella* planula [41] with our tissue-specific transcriptome; this revealed the cilia genes enriched in the apical tuft. Overall, we provided a comprehensive knowledge base of the larval sensory structure, instrumental for investigating the functional mechanism and signalling pathways associated with ciliated larval behaviour.

**Figure 2:**
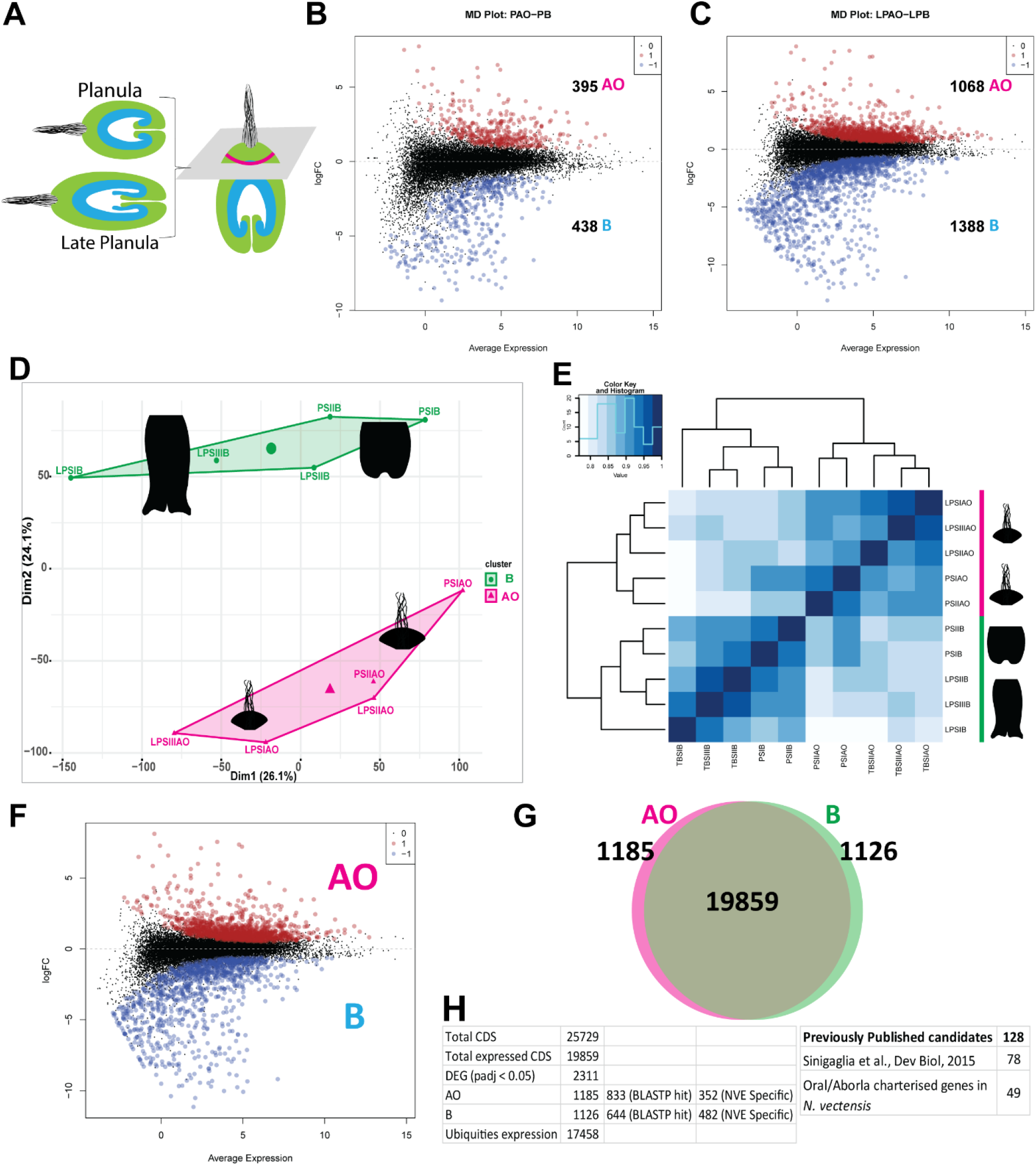
Global transcriptomic analysis. **(A)** Schematic representation of apical region microdissected from the rest of the larval body tissue for transcriptomics. **(B-C)** MD-plots represent the logFC ratio of differential expression between apical and body tissues from **(B)** planula and **(C)** late planula stages. The upregulated and downregulated genes are highlighted as red and blue circles, respectively. **(D)** PCA plot displaying a global overview of all data sets. PAO: planula apical organ; PB: planula body; LPAO: late planula apical organ; LPB: late planula body. **(E)** Correlation analysis identified two major clusters, as noted in the PCA plot. **(F)** MD-plots represent the logFC ratio of differential expression between apical and body tissues, data sets pooled from planula, and late planula development stages. **(G)** Venn diagram showing the DEG from apical and body tissues. **(H)** A table detailing the number of genes identified as DE in the current study, the number of genes with homologs, and genes identified in previous apical organ study by Sinigaglia et al. (2015) [2] and the number of genes previously shown to associated with in oral/aboral in *Nematostella* planula. For additional details, refer to **Supplementary Table 1**. AO: apical organ; B: body.

## Results and discussion

### Transcriptome profile of *Nematostella* apical sensory organ

We performed microdissections on *Nematostella* larvae and carefully separated the apical tissue from the rest of the larval body at the planula (∼ 50-60 hours post fertilisation) and late planula (∼ 75-85 hours post fertilisation) developmental stages **(Fig 2A)**. We acquired transcriptome data from both the apical tissue and the rest of the body separately to perform differential gene expression (DGE) analysis. The DGE analysis showed statistically significant variations among the apical and body tissue in both planula and late planula stages **(Fig 2B & C)**. The late planula stage **(Fig 2C)** presented a relatively large number of differentially expressed genes (DEG) in comparison to planula stage **(Fig 2B)**. Further, to characterise global gene expression patterns among the apical and body tissues from planula and late planula stages, we compared the transcriptomic profiles of all data sets using principal component analysis (PCA) and correlation analysis **(Fig 2D & E)**. The plots displayed a strong correlation among the replicates and a significant difference between the apical and body data sets from both planula and late planula stages. Notably, correlation analysis identified two major clusters and as illustrated in the PCA plot, the apical datasets from both planula and late planula form a single cluster. Likewise, the body datasets fall under a single cluster irrespective of their developmental stage **(Fig 2F)**. For the downstream analysis, we pooled both apical data sets from planula and late planula developmental stages, likewise, we pooled the body data sets. A DGE analysis was carried out among the apical and body datasets to identify the significantly DEGs (padj (FDR) < 0.05). We identified 2311 DEGs of which 1185 were enriched in the apical domain and 1126 were body-enriched **(Fig 2 G-H, Supplementary Table 1)**. This analysis revealed an unexpected complexity between apical and body tissues and indicates the specialisation of these tissues.

To validate our data by in situ hybridisation (ISH), we selected a set of newly identified apical pole enriched genes from the tissue-specific transcriptome data (**Fig 3A**). ISH showed that their expression is principally localised to the apical organ **(Fig 3B-O)**. Strikingly, two distinct expression profiles were observed: probes specific to *NVE14902, NVE4712* & *NVE8226* transcripts are localised around the apical cells (**Fig 3 B-D**), while probes specific to *NVE27, NVE10492, NVE8009, NVE2832* and *NVE2235* transcripts (**Fig 3**) were localised specifically within the apical cells. This pattern was also identified with other *Nematostella* apical organ genes by Sinigaglia et al. (2015) and termed as spot and ring [2]. Overall, our study has revealed the expression profile of the apical region, providing an opportunity to characterise the function of the larval sensory organ **(Fig 2H, Supplementary Table 1)**.

**Figure 3:**
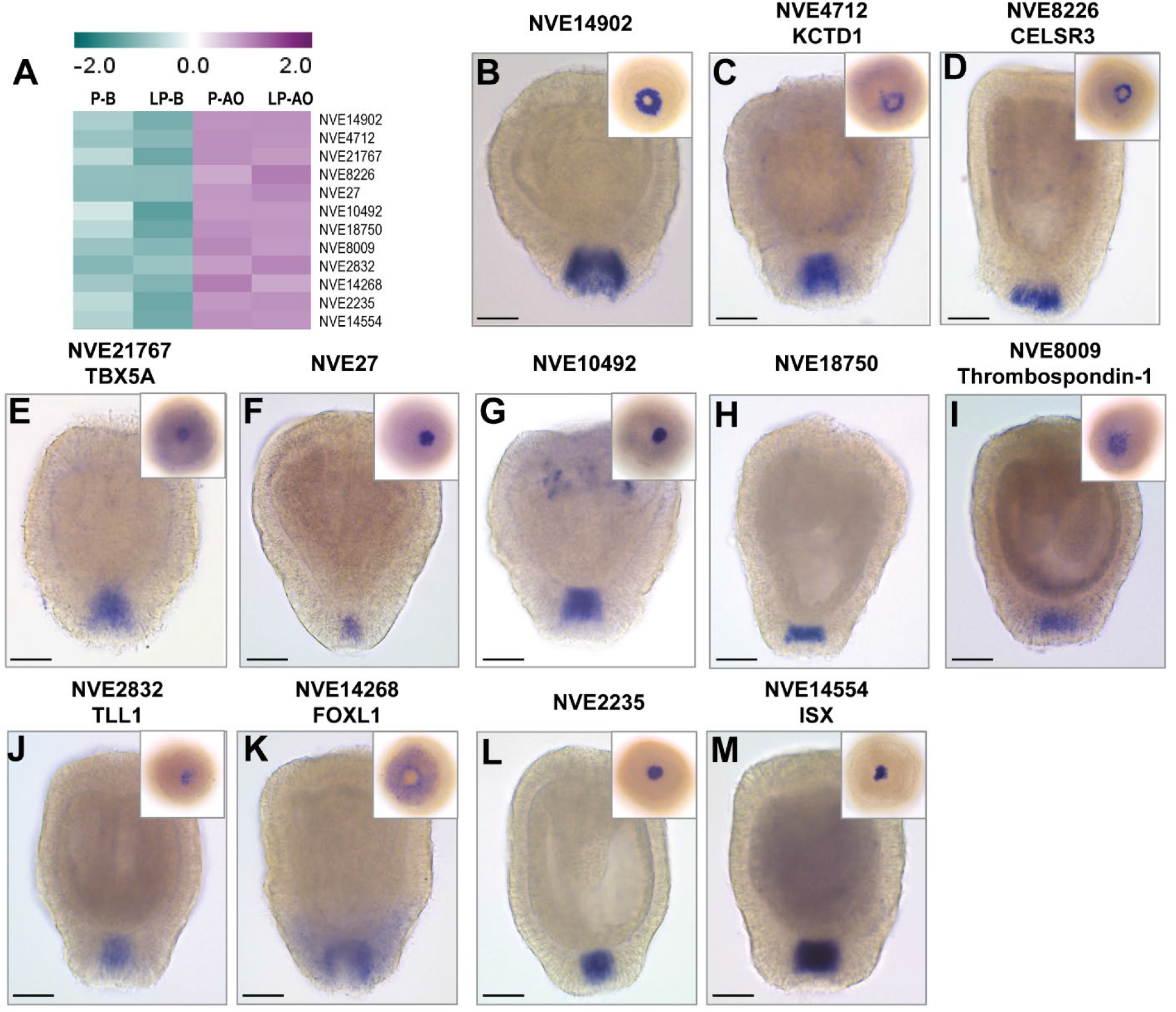
Validating apical tissue enriched genes. **(A)** A heatmap displaying the gene expression of selected marker genes enriched in apical organ cells. **(B-O)** ISH of apical organ enriched genes. At the inset on the right is the image displaying apical view. Scalebar = 50 µm

### Identifying the spatial distribution of *Nematostella* larval cell types by integrating tissue-specific transcriptomes with larval single-cell data

Fine morphological studies in different ciliated larvae [11, 42-44] have revealed that, along with long ciliated apical cells, the apical region comprises other cell types such as neurons, gland cells and peripheral ciliated sensory cells. Identification of apical organ associated cell types and their marker genes is strategic for understanding apical organ function. Single-cell RNA sequencing revealed transcriptome profiles of different cell types in ciliated larvae of *Nematostella* [40]; the larval cell types were classified into 38 metacells, and each metacell represents a specific larval cell type with a unique transcriptome profile [40]. However, the spatial distribution of these cell types has yet to be addressed. In our tissue-specific transcriptomes, the apical region displayed an expression profile distinctive from the rest of the body **(Fig 2)**. To develop an atlas of apical organ associated cell types and their transcriptomes, we integrated the tissue-specific transcriptome data with the *Nematostella* whole larval single-cell RNA-seq data [40].

We analysed the expression profiles of apical pole enriched genes (1185) in each larval metacell using cluster association (**Fig 4A**), PCA **(Fig 4B)**, and hierarchical clustering (HC) **(Fig 4C)** to define the metacells enriched in the apical domain. From the plots, we observed that the apical cell type is clustered distinctly from other cell types and had a high expression of multiple apical enriched genes **(Fig 4A)**. Strikingly, larva-specific neurons, gland/secretory cells 1 & 2, and undifferentiated cell types 2 & 4 also displayed specific expression of apical enriched genes **(Fig 4A)** and stood out in the apical domain with minimum overlap with any other larval cell clusters **(Fig 4A-C)**. In parallel, we also analysed the expression profile of the body enriched genes (1126) in each larval metacell to identify the cells enriched in body tissue **(Fig 4D-F)**. In both PCA and HC, genes enriched in the body displayed quite a distinctive trend from the apical tissue transcriptome **(Fig 4)**. We observed that the larval metacells including cnidocytes, gastrodermis, and a specific gland/secretory cell type 4 displayed high expression of body-enriched genes (**Fig 4D-F**) and clustered distinctly from other metacells **(Fig 4E & F)**, suggesting that these cell types are likely spatially distributed in the body and devoid from the apical region.

**Figure 4:**
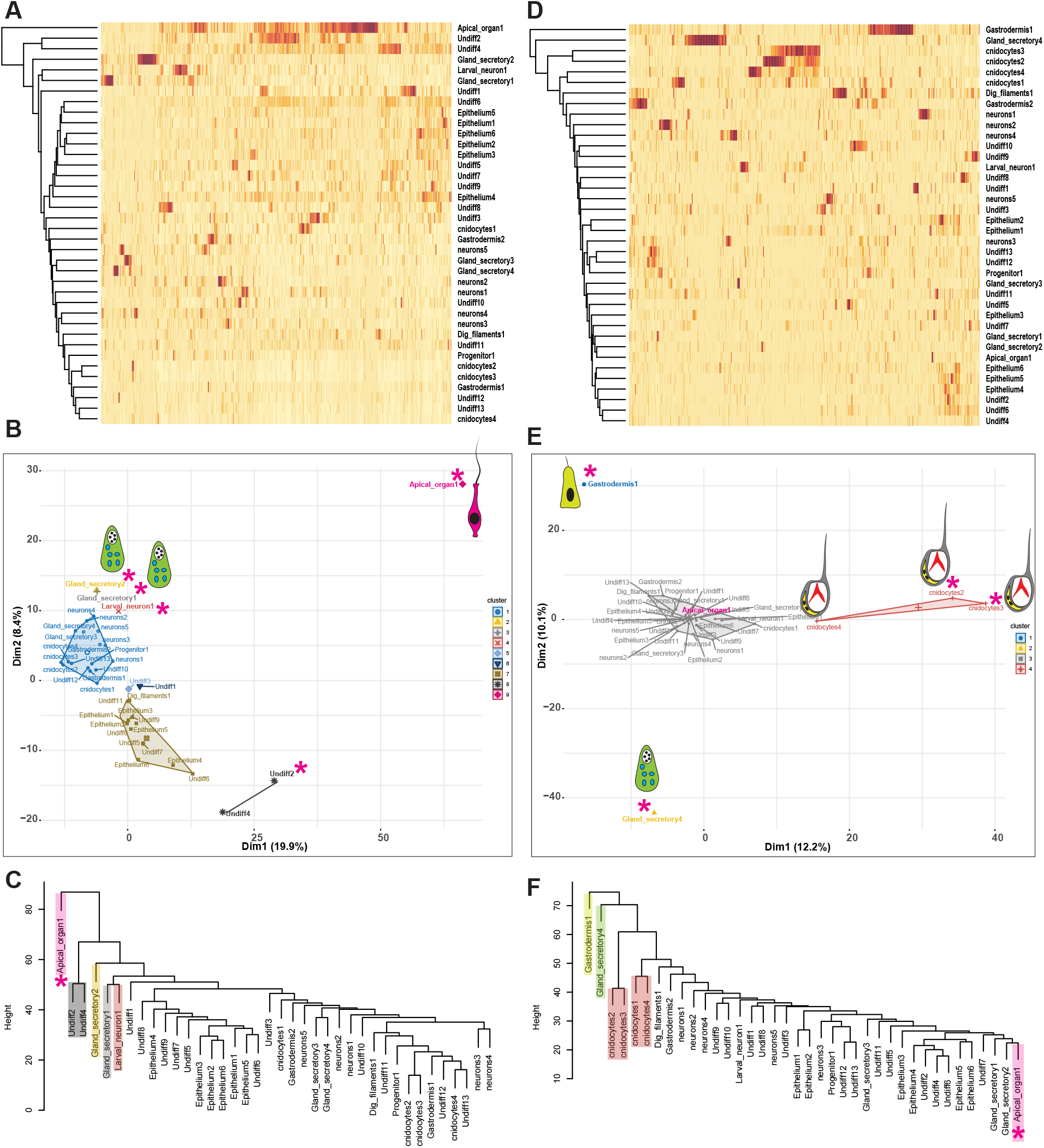
Spatial distribution of *Nematostella* larval cell types: Expression of apical **(A)** and body **(D)** enriched genes (rows) across 38 metacells sorted by cluster association. **(B & E)** Single-cell PCA plot for apical **(B)** and body enriched genes **(E). (C & F)** Hierarchical clustering, the dendrogram reproduced similar results as single-cell PCA.

Overall, this analysis clearly indicates the regionalisation of cell types in the larvae and provides an opportunity to characterise the function of the larval sensory organ in connection to other cell types potentially enriched in this area. Encouraged by these results, we set out to investigate the expression patterns of the cell-type marker genes in the planula through ISH to define the spatial distribution of larval cell types in the apical domain.

### *Nematostella* apical organ is composed of apical tuft cells and crowned with larval neurons

Based on the *Nematostella* single-cell transcriptome study [46], two metacells were classified as larval specific with low similarity to any adult cell cluster. Based on previously identified marker genes *Fgf1a* [2] and *Nk3*, one of these metacells was recognised as apical cell type. The second metacell showed a peculiar expression of genes with putative neuronal functions such as cyclic nucleotide-gated, TrpA, polycystic kidney disease, and shaker ion channels therefore, it was classified as an uncharacterised larval-specific neuronal cell type [40] which remained to be characterised.

From single-cell PCA and HC analyses, we observed that along with the apical cell type, the larva-specific neurons stood out in the apical domain **(Fig 4A-C)**. In our initial ISH analysis of apical enriched genes, we observed two distinct expression profiles: a set of genes expressed around the apical cells **(Fig 3 B-D)** and others expressed specifically in apical cells (**Fig 3 B-D**). The distinct expression patterns of apical organ genes suggest that apical sensory structure is composed of two distinct cell types. Based on the *Nematostella* single-cell transcriptome [40], the marker genes identified as spot-specific were principally restricted to the apical cell type. On the other hand, the marker genes expressed as a ring were enriched in larval neurons **(Fig 5)**.To explore this further, we carried out double FISH on these marker genes **(Fig 5)**. To our surprise, *NVE8226* & *NVE14902* markers, besides being expressed as a ring around the apical cells, showed expression across the whole apical region **(Fig 5A & E; Movie 1 & 2)**. The cells have a peculiar sensory neuronal morphology [20] and are localised in the ectoderm with the tip pointing towards the peripheral direction **(Fig 5B & F)**. Expression of these genes gradually diminishes as the larvae progress through metamorphosis **(Fig 5C, D, G & H)**, reflecting their larval specific function. Based on the double FISH, the apical organ gene *Alx* (*NVE14554*) is restricted to the apical pit and crowned with larva-specific neurons (detected by *NVE8226* & *NVE14902* markers) **(Fig 5J-M; Movie 3 & 4)**. The cells with apical tuft were concentrated in the apical pit visualised by immunostaining with an anti-tubulin antibody, where the spot genes like *Alx* are expressed **(Fig 5N)**. The current analysis allowed us to localise the previously uncharacterised larval-specific neuronal cell type to the apical organ ring. Such localisation suggests that the apical cells with their ciliated tuft act as a sensory structure, and the larva-specific neurons receive information by crowning around them and probably signalling downstream to the rest of the body **(Fig 5O)**. This postulation coincides with the previous finding from the *Nematostella* single-cell sequencing study [40]. The apical organ cells lack any recognisable neuronal effector genes such as G protein coupled receptors (GPCRs), synaptic scaffold proteins, neurotransmitter-related enzymes, and neuropeptides, suggesting a non-neuronal identity [40]. On the other hand, the larval-specific neuronal cell type is enriched with neuronal functional genes. To conclude, the apical tuft cells and larval-specific neurons together form an active apical sensory organ **(Fig 5O)**.

**Figure 5:**
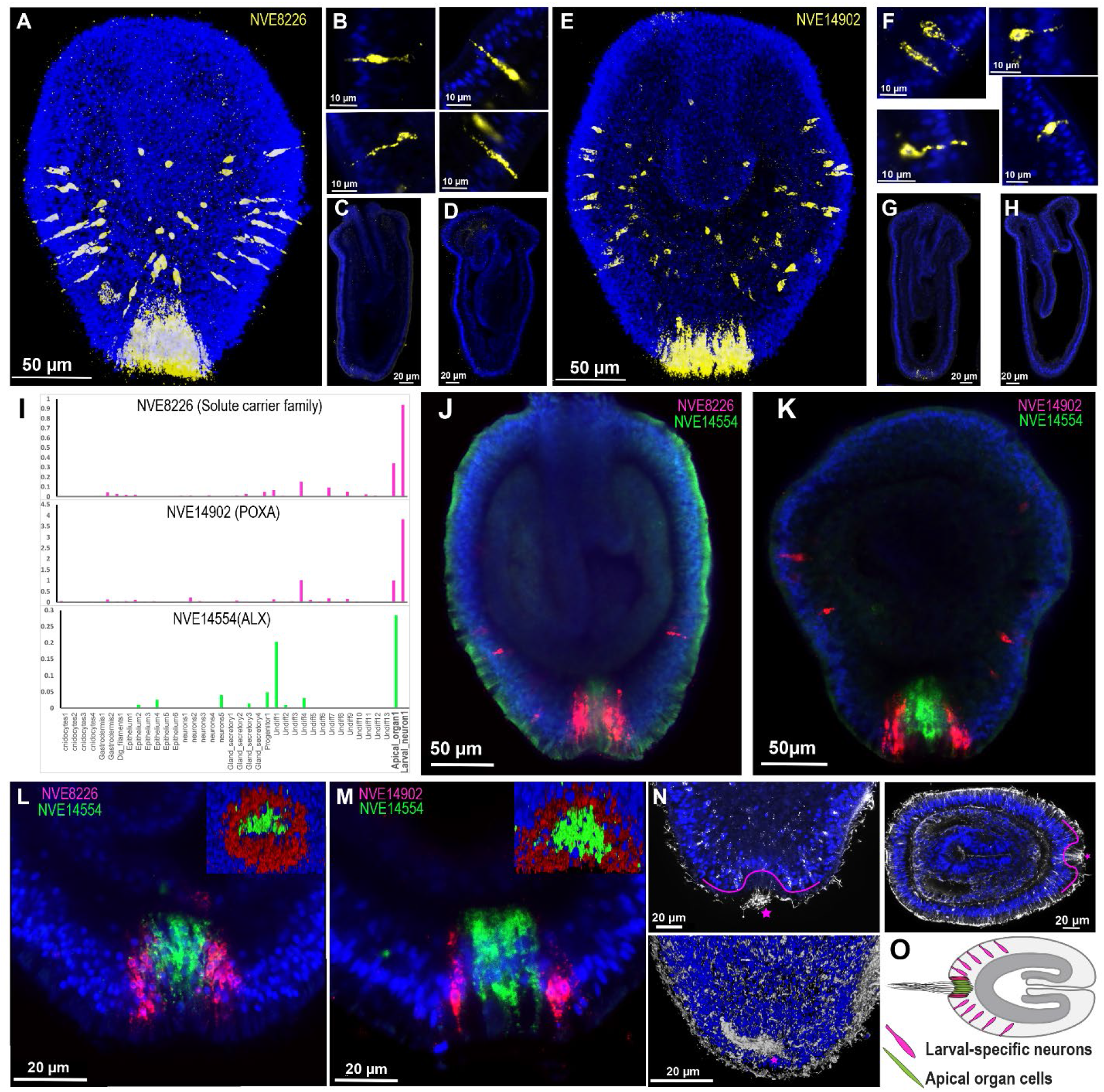
Whole-mount fluorescent in-situ hybridization (FISH) staining of the apical organ and larval-specific neuron genes. (A-D) Expression of *NVE8226* marker gene (yellow). (E-H) Expression of *NVE14902* marker gene (yellow). (I) The bar plot displays the expression profile of selected marker genes. (J-M) Co-localisation of marker genes for the spot *(NVE14554, Alx*; green) and the ring (*NVE8226* & *NVE14902*; red) by double FISH. DAPI is shown in blue. (L & M) The inset on the left displaying the 3D image from apical view. (N) Immunostaining (white) with the anti-acetylated tubulin antibody counterstained with DAPI (blue) for nuclei; the pink line demonstrates the apical tuft cells concentrated in the apical pit. (O) Summary diagram showing the spatial distribution of apical organ/tuft and larval-specific neuronal cell types.

### Flask-shaped gland/secretory cells enriched in the apical domain

Besides apical cells and larva-specific neuronal cell types, the single-cell PCA analyses revealed four gland/secretory cell types with distinctive profiles among apical and body data sets **(Fig 4A-D)**. Out of four larval gland cell types, cell types 1 and 2 are principally enriched in the apical region **(Fig 4A&C)**. In contrast, gland/secretory cell type 4 is enriched exclusively in the body **(Fig 4B&D)**, while cell type 3 expressed ubiquitously **(Fig 4A-D)**. Recent studies have addressed the development of gland cells in *Nematostella*, mainly focusing on gland cells that develop and integrate into the pharynx and mesenteries of polyps [45, 46] and ectodermal gland cells producing toxins [47, 48]. However, not much focus has been given to other larval gland/secretory cell types and their fate during development. To confirm the enrichment of gland/secretory cell types in the apical domain, we selected specific marker genes expressed in each of these gland cell types and visualised their expression pattern by ISH **(Fig 6)**. Gland cells type 1 and 2 were enriched in the apical region **(Fig 6A-F; Movie 5)**. Gland cell type 3 is presented throughout the animal **(Fig 6G, I & J; Movie 6)**. The marker genes (*NVE23810, NVE26086, NVE9234* & *NVE10584*) selected for gland cell type 4 are expressed in the pharynx region and progress into the pharyngeal/mesentery tissue of the primary polyp **(Fig 6K-O)**. In contrast, marker genes for cell types 1, 2 and 3 were expressed in the outer ectoderm of the planula **(Fig 6A-J)**.

**Figure 6:**
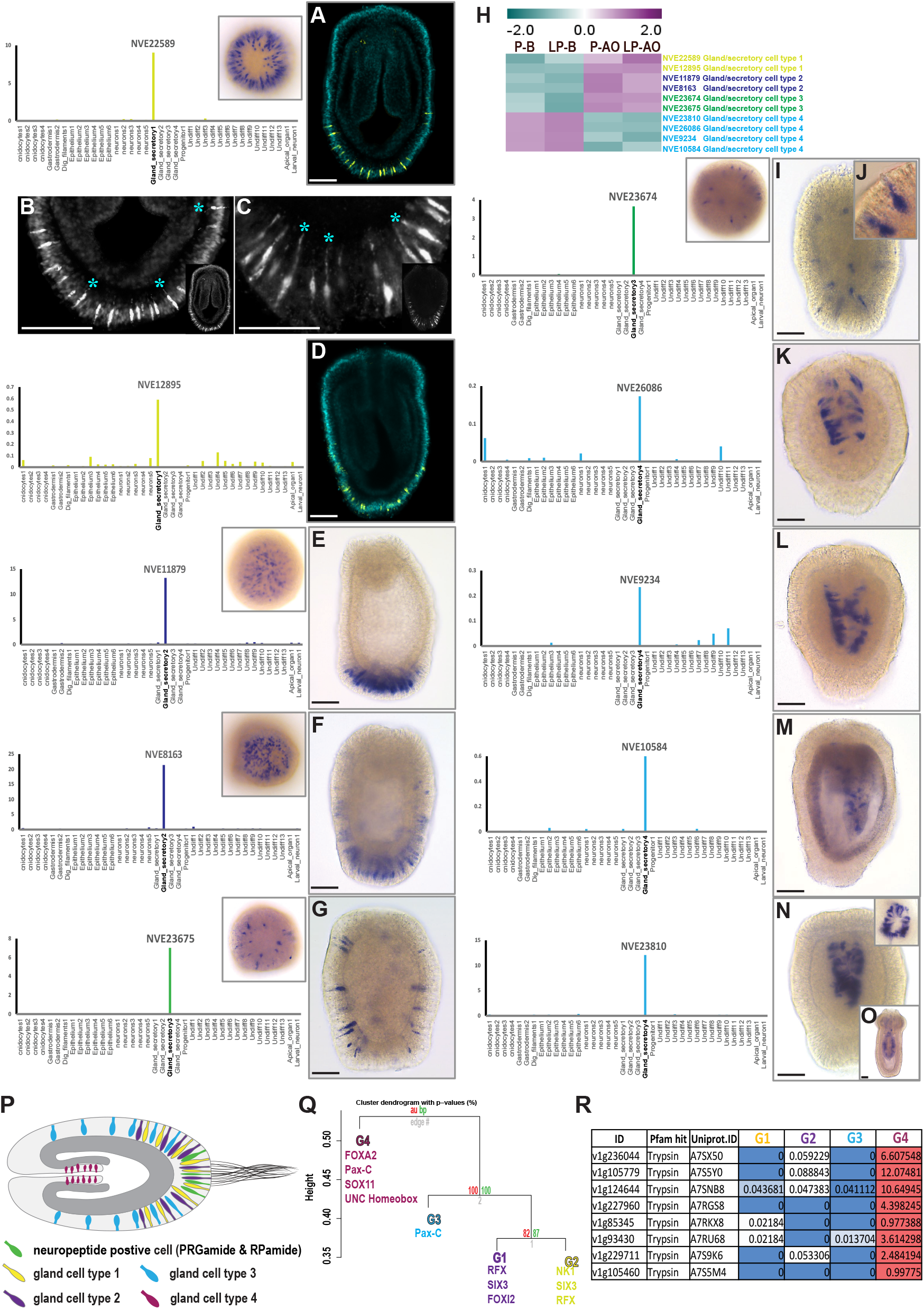
Spatial distribution of larval gland/secretory cell types visualised by whole-mount ISH. **(A, D-G, I-O)** (Right) ISH of larval gland/secretory cell type marker genes, (Left) each bar plot displays the expression profile of selected marker genes in the different larval cell populations. At the inset on the right is the image displaying an apical view. **(A-D)** ISH of larval gland/secretory cell type 1. **(B-C)** Monopolar sensory cells with their projection from the cell body extended towards mesoglea (*), the inset showing the whole animal. **(E & F)** ISH of larval gland/secretory cell type 2. **(G, I & J)** ISH of the larval gland/secretory cell type 3, panel H shows the ciliated gland cell type 3 at higher magnification. **(H)** A heatmap displays the gene expression pattern of selected marker genes specific for each larval gland/secretory cell types. **(K-O)** ISH of the larval gland/secretory cell type 4, panel P demonstrates the spatial expression of gland cell type 4 concentrated in the mesentery tissue. **(P)** A schematic representation of larval gland/secretory cell type’s spatial distribution. **(Q)** A dendrogram displaying the correlation among gland cells and a list of a different combination of transcription factors expressed in each gland cell type. **(R)** Gene expression of trypsin domain-containing proteins in gland cell of larvae single-cell data. Scalebar = 50 µm.

To gain further insights into molecular features of the larval gland cell types, we looked into the *Nematostella* larvae single-cell data [40]. We noted that each of these gland cells are regulated through a set of transcription factors and some are unique to specific gland cell types **(Fig 6Q)**. For instance, gland cell 1 expresses homeobox-OAR (*v1g184843*) and FOXI2 (*v1g96685*) and gland cell 2 expresses NK1 transcription factor (*v1g8907*). On the other hand, some of these transcription factors are commonly expressed between two glands cells, such as homeobox-PAX (*v1g168908*), expressed in both gland cell 3 and 4. Similarly, the RFX4 (*v1g122918*) is expressed in both gland cells 1 and 2 **(Fig 6Q)**. This suggests that a different combination of transcription factors modulates the trajectory of each of these gland cells. As noted from ISH, gland cell 4 has a distinct expression from other peripheral gland cells and is principally restricted to pharyngeal/mesentery tissue **(Fig 6K-O)**. From the *Nematostella* larvae single-cell data, we noted that the trypsin domain-containing proteins are restricted to gland cell 4 **(Fig 6R)**, suggesting that gland cell 4 are the larval cell type that develops into polyp gland/secretory cell types in the mesenteries, where the digestive enzymes like trypsin are synthesized and secreted into the gastrovascular cavity for digestive function [46].

### Spatial distribution of aboral enriched neuropeptide expressing cells and gland cells

Besides larval specific neurons and gland cells, we also noted a significant difference in the spatial distribution of neuropeptide transcripts along the oral-aboral axis in *Nematostella* **(Fig 8A)**. Along with the previously identified Nv-RPamide III neuropeptide [22], we also detected PRGamide is exclusively expressed in the apical tissue **(Fig 7A, B&C)**, while Nv-LWamide, Antho-RFamide, neuropeptides type 2 and HIRamide were detected predominantly in the body tissue **(Fig 7A, D&E)**.

**Figure 7:**
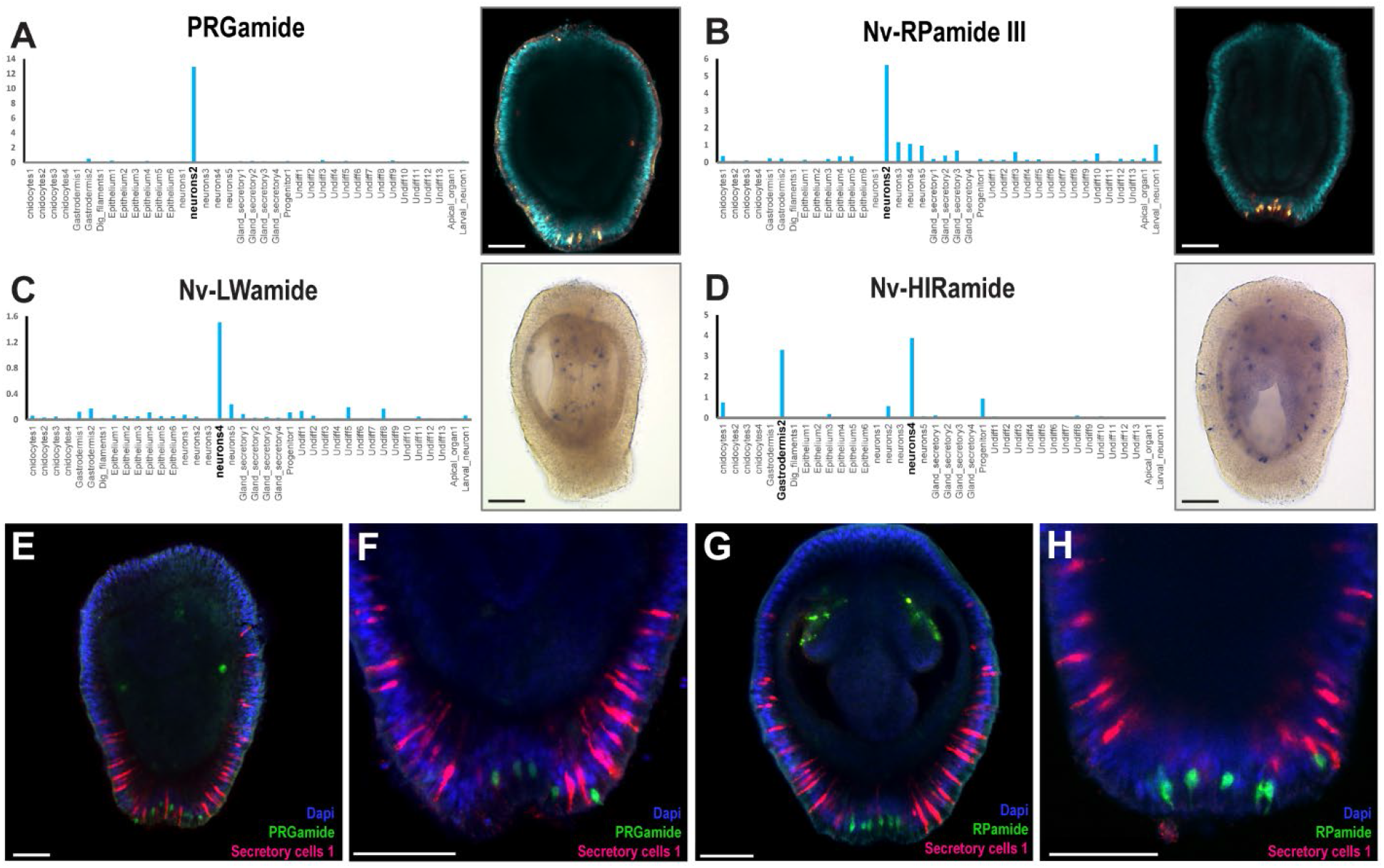
Spatial distribution of neuropeptide expressing cells. **(A-D)** ISH of selected neuropeptide genes, each bar plot displays the expression profile of selected marker genes in a different larval cell population (left). **(A&B)** Neuropeptides Nv-RPamide III and PRGamide were exclusively expressed in the apical tissue **(C&D)**, while Nv-LWamide and HIRamide were detected predominantly in the body tissue. **(E-H)** Double fluorescence in situ hybridisation for gland/secretory cells type 1 (magenta) and either PRGamide or Nv-RPamide III (green). DAPI is shown in blue. Scalebar = 50 µm.

Overall, we discovered that the apical domain comprises a minimum of 6 distinct cell types, including apical cells, neurons (larval specific neurons, neuron type 2), gland cell type 1 and 2 and undifferentiated cell types 2 and 4. Besides apical cells and larval-specific neuronal cell types, the distinct expression of the gland cell types 1 and 2 and neuropeptide positive cells (Nv-RPamide and PRGamide) in the apical region appears intriguing. To explore this further, we carried out double FISH on marker genes of the apical enriched gland/secretory cell type 1 and neuropeptides PRGamide and Nv-RPamide III **(Fig 7E-H)** to visualise their relative spatial distribution in the apical region. Notably, the gland/secretory cells of type 1 are large in number with a broad distribution relative to neuropeptide expressing cells. We also noted that the gland cells exhibited a typical flask-shaped morphology. Such peripheral flask-shaped cells with similar spatial distribution are also identified in a range of marine phyletic taxa including sponge [27, 28, 49, 50] and eumetazoan larvae [12-14], signifying a deeply conserved larval sensory/secretory system. Hypothetically, the aboral peripheral secretory mechanisms predated the Eumetazoa, suggesting that the ancient sensory/secretory mechanisms already evolved prior to neurons and gradually progressed into a diffused sensory system interspersed among the motile ciliated cells of the epithelial layer (sensory and ganglion neurons). Based on the morphological similarities and their spatio-temporal expression **(Fig 6A-D, Fig 7F-I)**, we speculate that the peripheral gland/secretory cells of *Nematostella* may have a crucial role in metamorphosis. Further studies in this direction may elucidate the functional relationship among gland/secretory cell type 1 with neurons and apical cells.

### A distinct neuronal regulatory network associated with the larval apical region

The DE of neuropeptides along the larval body axis **(Fig 8A, D&E)**, suggests that neuropeptide networks differ between the apical and body tissue [19]. To gain further understanding of the neuronal mechanisms linked with the apical organ, we explored the tissue-specific transcriptomes using gene ontology (GO) to identify genes related to neurotransmission gene networks. We performed a BLASTP search in the Uniprot database to identify the protein homologs. Additionally, we used gene functional annotation data from the published *Nematostella* single-cell transcriptome study [40] **(Supplementary Table 1)**. Next, we analysed their gene ontology terms using the David 6.7 and PANTHER 15.0 gene ontology tools. We identified several neuronal associated genes that are DE between the apical and body regions, such as neuropeptides, G-protein coupled receptors, ligand-gated ion channels and neurotransmitter synthesis genes (**Fig 8A**) **(Supplementary fig 1)**. This indicates that the neuromodulation in the apical region differs from the rest of the body and is characterised by a specific set of genes in *Nematostella*. Each of these genes may potentially be involved in modulating larval swimming behaviour. For example, in annelid and echinoderm larvae, the ciliary beat frequency is modulated by neurotransmitters, and neuropeptides can actively control the swimming speed by modulating the ciliary beating frequency [5, 6, 51, 52].

**Figure 8:**
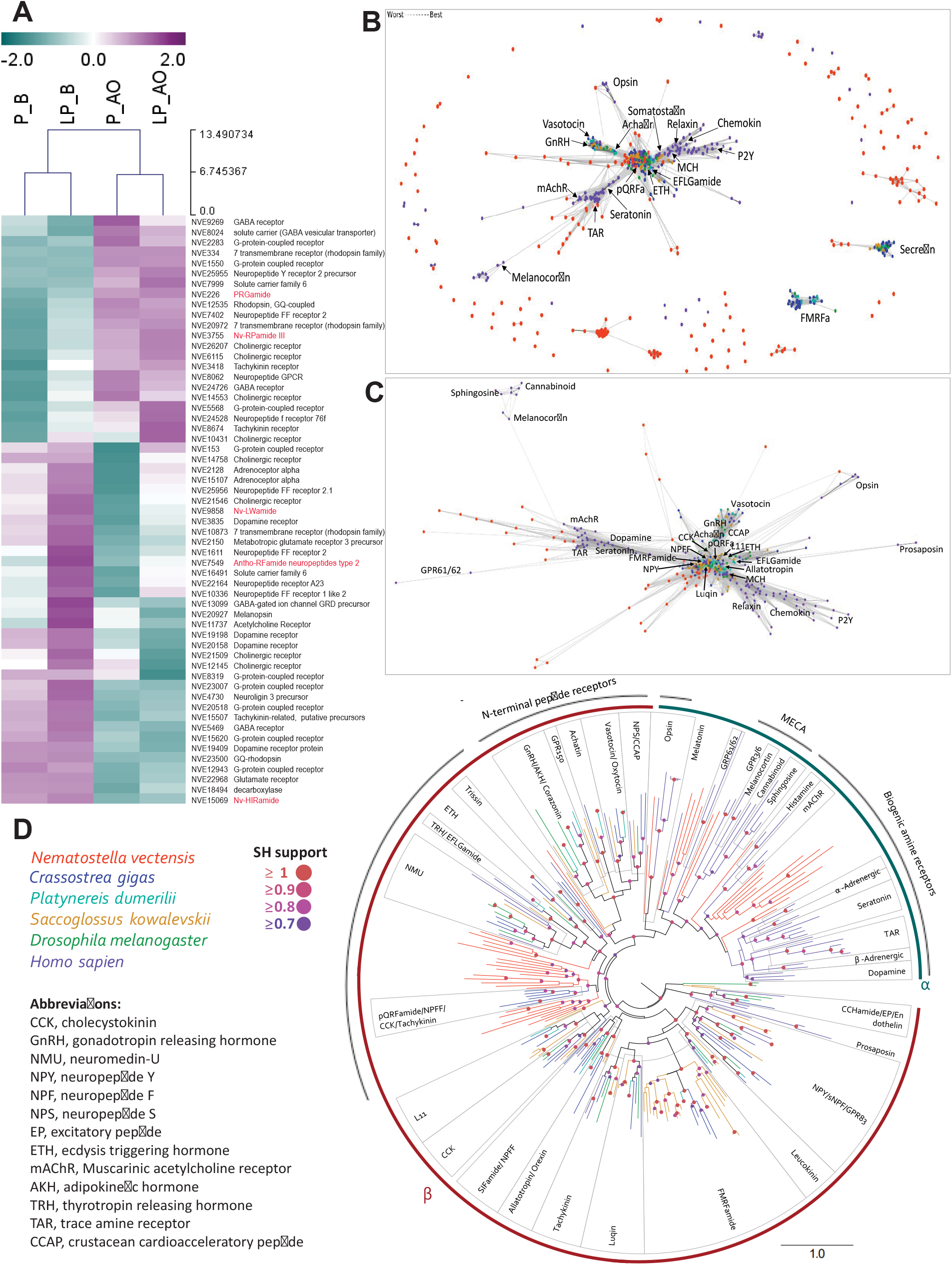
Spatial distribution of the larval nervous system associated genes. **(A)** A heatmap displaying the gene expression pattern of a set of genes related to diverse neuronal functions. **(B)** Sequence-similarity-based clustering of all *Nematostella* apical enriched GPCRs. Forty-six *Nematostella* GPCRs clustered with a large group of Rhodopsin and four clustered with Secretin. **(C)** Cluster map of the Rhodopsin cluster presented in B. **(D)** Maximum likelihood analyses of the sequences from cluster analysis C. Phylogeny constructed with *Crassostrea gigas, Platynereis dumerilii, Saccoglossus kovalevski, Drosophila melanogaster* and *Homo sapiens*.

The neurotransmitters were demonstrated to control the ciliary beating through two broad classes of receptors: ligand-gated ion channels and GPCRs [7, 53] [51, 54]. In echinoderms, the ciliated larvae actively respond to contact with obstacles [55, 56] and this action is potentially mediated by ionotropic acetylcholine receptors [55, 57-59]. The expression profiles of transcripts coding for ligand-gated ion channels such as neuronal acetylcholine receptor, glycine receptor and beta subunit of the gamma-aminobutyric acid receptor are segregated among apical and body tissue in *Nematostella* **(Supplementary Fig 1)**. Further, genes associated with pre/post-synaptic neurons and neurites conducting nerve impulses such as synaptotagmin, neural cell adhesion molecule, tyrosine kinase-like orphan receptor, sodium-dependent dopamine transporter, guanylate cyclase soluble subunit, beta-1 glutamate receptor, calmodulin, vesicular glutamate transporter and netrin were DE among oral and apical tissue **(Supplementary Fig 1)**.

We observed a large number of GPCRs DE among oral and apical tissues **(Supplementary Fig 1)**. GPCRs are known for their role in neuropeptide and other neurotransmitters signalling [60]. To gain further understanding of the relationship of apical enriched GPCRs with known GPCR families in Bilateria, we performed sequence-similarity-based clustering and Maximum likelihood phylogenetic analyses on apical enriched GPCRs, in parallel to BLASTP analysis.

We used previously published datasets from *Crassostrea gigas, Platynereis dumerilii, Saccoglossus kovalevski* and *Drosophila melanogaster* [61], as well as *Homo sapiens* GPCRs [62]. The *Nematostella* apical enriched GPCRs clustered with a range of GPCR superfamilies from bilaterians as shown in Figure 7B, C & D. Forty-six *Nematostella* GPCRs clustered with a large group of rhodopsin and four clustered with secretin (**Fig 8B**). Within the rhodopsin superfamily, the GPCRs formed two major groups: rhodopsin α and β. Rhodopsin α GPCRs formed a monophyletic group which included twenty two *Nematostella* GPCRs. The remaining twenty four *Nematostella* GPCRs fell among the rhodopsin β clades (**Fig 8D**). The rhodopsin α GPCRs can be further divided into biogenic amine receptors which contained thirteen *Nematostella* GPCRs, most of which formed a sister clade with histamine and muscarinic acetylcholine receptors from humans. The remaining *Nematostella* GPCRs within rhodopsin α clustered with the MECA group that includes melanocortin and opsins. Among the rhodopsin β GPCRs twenty three *Nematostella* GPCRs formed three clades positioned among other neuropeptide GPCRs within the rhodopsin β superfamily. Four *Nematostella* GPCRs formed a clade that shares a common ancestor with bilaterian NMU, TRH, EFGLamide, ETH and Trissin. Forming a sister group to this were twelve *Nematostella* GPCRs that were orthologous to a set GPCRs including pQRF, NPFF, CCK and tachykinin receptors from the mollusc *Crassostrea* (pQRF, CCK and tachykinin receptors from *Crassostrea* also occurred elsewhere within the phylogeny, clustering with genes of similar function from other bilaterians). In summary, the current analysis allowed us to identify several ortholog groups of GPCRs expressed in the *Nematostella* apical domain and their distribution across different GPCR receptor families.

Overall, these observations indicate that the molecular identity of the apical and body tissue is determined by the specific expression of neurotransmitter receptors, ion channels and neuropeptides. Thus, given the complexity and vast differences in nervous system organisations between the apical and body domains, further studies in this direction can elucidate the differences between the larval and adult nervous systems. This will reveal what neuronal mechanisms are active in ciliated larvae and how the larval specific nervous system is utilised to modulate swimming behaviour.

### The molecular composition of apical tuft cilia

The body-enriched genes revealed GO terms such as extracellular matrix organisation, cell differentiation, nervous system development and neuron differentiation. In contrast, the apical domain enriched genes revealed GO terms principally associated with the cilium, embryonic morphogenesis, photoreceptors and synaptic signalling **(Fig 9A & B)**. Among the cilium related GO terms enriched in the apical region (cilium organisation, photoreceptor connecting cilium, and motile cilium) we identified a set of genes associated with a ciliary membrane and axonemal-associated genes (intraflagellar transport (*ift*), *bardet-biedl syndrome (bbs)*, kinesin family member (*kif*), enkur, ADP ribosylation factor like GTPase (*arl)*, filamin A (*flna*), tetratricopeptide repeat domain (*ttc*), regulatory factor X3 (*rfx3*), stabiliser of axonemal microtubules 1 (*saxo1*), usherin (*ush*), tektin (*tekt*), β-tubulin (*tbb*) & tubby bipartite transcription factor (*tub*)).

**Figure 9:**
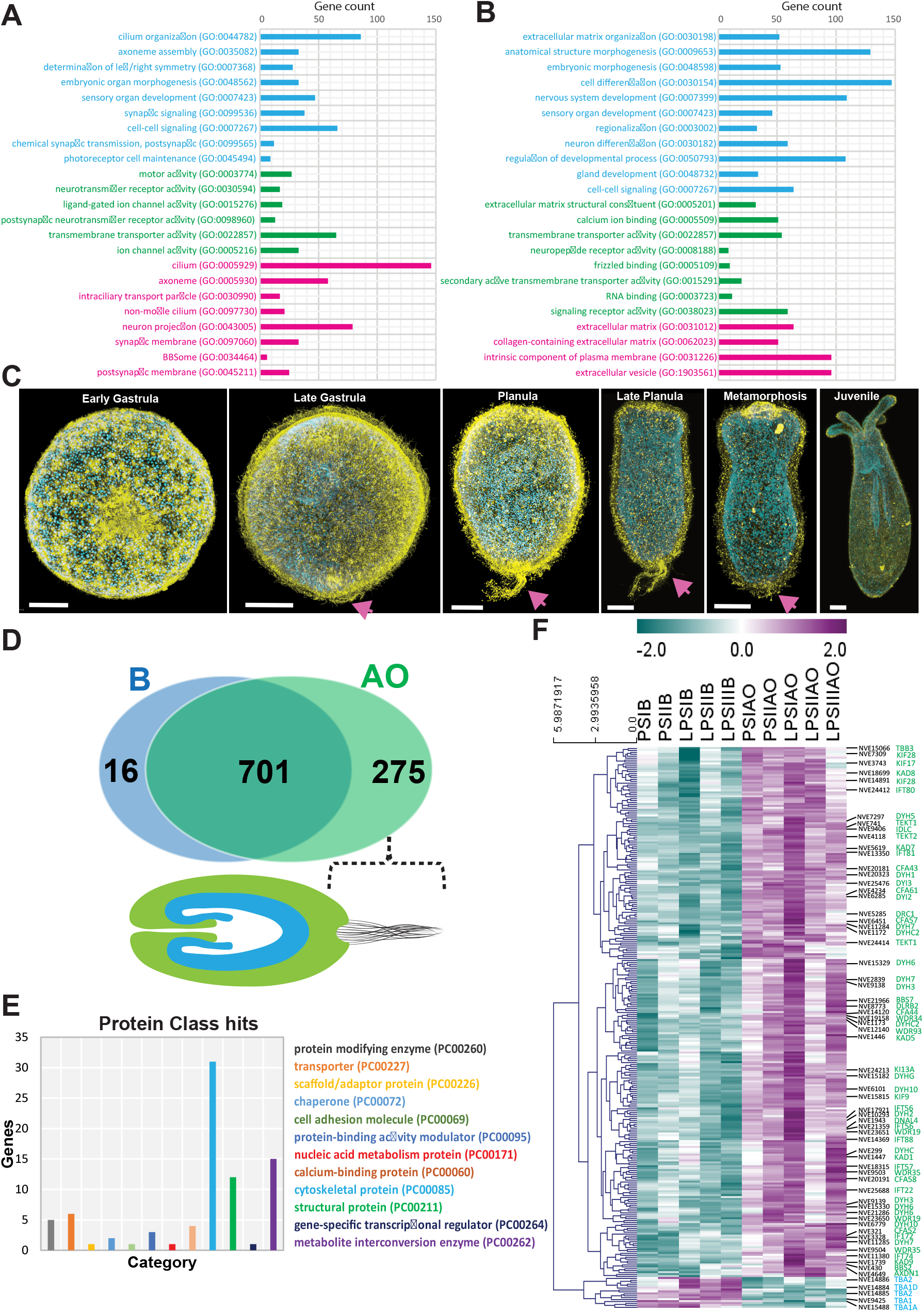
Gene ontology analysis of significantly differentially expressed genes. **(A)** Enriched GO terms among apical specific genes. **(B)** Enriched GO terms among body-specific genes. **(A & B)** GO category includes biological processes (blue), molecular function (green) and cellular components (pink). **(C)** Different stages of *Nematostella* development. Immunostaining (yellow) with the acetylated tubulin antibody and counterstained with DAPI (blue) for nuclei. Arrow pointing to apical tuft. **(D&E)** Tissue-specific transcriptome vs ciliary proteome of *Nematostella* planula to define cilium associated genes enriched in apical turf. **(D)** Venn diagram showing the DEG from apical and body tissues. **(E)** Cilia genes categorised into different protein class, the top hits against cytoskeleton category (31 genes). **(F)** Heatmap displays the DE of genes identified in cilium proteome and differentially expressed between the oral and apical region of *Nematostella* larvae. Highlighted candidates associated with cilia assembly and maintenance such as intraflagellar transport protein (IFT), dynein heavy chain, axonemal (DHY), beta-tubulin (TBB), tektin (TEKT). Expression levels are normalised by Z-score. Scalebar = 50 µm.

The enrichment of ciliary related genes in the apical tissue is probably linked with the apical tuft **(Fig 9C)**. The apical tuft is built of non-motile cilia and found in a wide range of marine invertebrates such as echinoderms [63], molluscs [64], annelids [16, 65] and anthozoans. In contrast, motile cilia typically cover the entire body of the larvae and aid in swimming [63]. In general, cilia are involved in a diverse range of biological tasks, mainly sensory functions such as detecting environmental cues [66, 67]. Ciliated larvae have both motile and non-motile cilia [1] and the functional role of apical tuft cilia has largely remained enigmatic. However, its strategic position and its association with dense neurons (peptidergic neurons) downstream suggests that the apical tuft is a part of the sensory organ.

A comparative study of the ciliary proteomes in *Nematostella* planulae along with sea urchins and choanoflagellates [41] has revealed core components of the ciliary intercellular signalling pathways and identified the shared ciliary proteome. In that study, ciliary proteome data were acquired by isolating cilia from whole *Nematostella* planula, including the apical tuft, which were subjected to mass spectrometry; this allowed construction of the ciliary proteome from the whole larvae, including the apical tuft [41]. To reveal the functional role of the apical tuft, we characterised the difference in molecular composition between the apical tuft cilia and regular motile cilia.

We integrated our tissue-specific transcriptomes with cilia proteomes from *Nematostella* planula, which revealed an unexpectedly complex relation between apical tuft and motile cilia. As shown in **Figure 9 D&E**, among the DE genes, 275 ciliary genes were enriched in the apical region, while 701 ciliary genes were commonly expressed all around the body **(Figure 9 D&E, Supplementary Table 1)**. Thus, the apical tuft cilia possess a set of proteins distinct from the rest of the body cilia. A pioneer study in *Nematostella* apical organ by Sinigaglia et al. have identified 52 putative cilia-related genes enriched in the apical organ [2] and among these 26 are principally associated with cilia assembly and maintenance. In line with previously identified cilia genes, here we revealed 275 ciliary genes enriched in the apical region, likely associated with apical tuft.

Among the apical enriched genes, we came across a set of candidates related to cytoskeletal and structural proteins, such as dynein heavy chain, axonemal (DHY), beta-tubulin (TBB), tektin (TEKT) and intraflagellar transport protein (IFT) **(Fig 9E&F)**. Among the body enriched candidates, we identified a set of tubulin genes such as TBA1 (NVE9425), TBA1A (NVE15488), TBA2 (NVE14886), TBA1D (NVE14884), and TBA2 (NVE14885), likely this set of genes specifically aid in motile cilia assembly **(Fig 9F)**. Next, we observed genes associated with metabolite interconversion such as WD repeat-containing protein and kinase family members (phosphoenolpyruvate carboxy kinase, nucleoside diphosphate kinase, adenylate kinase) **(Fig 9F)**. In summary, the current analysis highlighted the composition of the apical tuft and provided a prime resource for further studies to characterise the function of the apical tuft.

### Conclusions

We revealed a comprehensive molecular signature of the *Nematostella* sensory structure. Using transcriptomics and ISH, we unravelled the cell types enriched in the larval apical region: apical cells, neurons, and gland/secretory cells. Strikingly, among the four types of larval gland cells, we identified two types associated explicitly with the apical sensory structure. The GO enrichment analysis revealed a considerable difference between the apical and body regions. The GO terms suggest that the apical organ is likely involved in sensory function through cilia with a unique neuronal network. The present study provides new insights into the apical organ signalling pathways for future mechanistic studies and genes associated with oral/aboral differentiation. Additionally, this resource identifies numerous candidate genes for elucidating the functional role of the apical organ.

## Methods

### *Nematostella* culture

*Nematostella* polyps were grown in 16 ‰ artificial seawater at 18°C in the dark and fed with freshly hatched *Artemia* nauplii. The induction of spawning was performed as previously described [68]. After fertilisation, the gelatinous substance around the eggs was removed using 4% L-Cysteine (Sigma-Aldrich, USA) [68].

### Microdissection of *Nematostella* apical organs

We performed microdissection on *Nematostella* larvae to separate the apical organ from the rest of the larval body. Two developmental stages including planula and late planula were used. The apical organ was isolated from the whole larvae using a 34 gauge needles and a stereomicroscope with 10X magnification, the area of the apical tissue defined in **Figure 2A**. Each sample was pooled from a minimum of 50 individual larvae and also included samples from a minimum of three different batches. The samples were carefully collected using glass pasteur pipettes and excess media was removed before snap freezing in liquid nitrogen and stored at -80°C until further processing.

### RNA sequencing and differential gene expression

For RNA isolation, due to the sheer size samples collected from multiple batches were combined to acquire an adequate amount of RNA for sequencing. Total RNA was isolated using the TRI Reagent® according to the manufacturer’s protocol. RNA quality was assessed using Agilent RNA 6000 Nano Kit on Agilent 2100 Bioanalyzer (Agilent, USA), and samples with RNA integrity number ≥ 8.0 were used for sequencing. The SENSE mRNA-Seq Library Prep Kit (Lexogen GmbH) was used for library preparation. Before sequencing, the libraries were pre-assessed by Agilent High Sensitivity DNA Kit (Agilent, USA) and quantified using Qubit™ 1X dsDNA HS Assay Kit (Invitrogen™). The sequencing was outsourced (GENEWIZ Illumina NovaSeq/HiSeq 2×150 bp sequencing), generating 20 million paired-end reads per replicate. Raw data were deposited at NCBI GEO submission GSE159166. After de-multiplexing and filtering high-quality sequencing reads, the adapter contamination was removed by using Trimmomatic v0.36 [69]. The quality of the reads was verified using FastQC [70]. Processed reads from each sample were mapped to the *Nematostella* genome (indexed bowtie2 [71]) by using STAR [50]. The number of reads mapping to each *Nematostella* gene models (https://figshare.com/articles/Nematostella_vectensis_transcriptome_and_gene_models_v2_0/807696) were extracted using HTSeq-count v0.6 [51]. Differential expression analyses were performed using limma (Galaxy Version 2.11.40.6) [72] and DESeq2 (Galaxy Version 2.11.40.6) [72]. The principal component analysis (PCA), hierarchical clustering (HC), and heat maps were generated using the R package in R-studio (Version 1.2.5019). For functional annotation, we used blastp [73] with the default curated gathering threshold to predict the protein homologs against the Uniprot database. Additionally, we used gene functional annotation data from a published *Nematostella* single-cell transcriptome study [40]. The Gene Ontology (GO) term enrichment was performed using gene annotation tools including PANTHER™ Classification System [74] and DAVID [75].

### In situ hybridisation (ISH) and double fluorescent ISH (dFISH)

ISH was performed according to published protocols [76, 77]. In brief, fixed animals were transferred into sieves and rehydrated in 1 mL 60% methanol/40% PBST and then washed in 30% methanol/70% PBST. Samples were digested in proteinase K (80 µg/mL) for 5 min then blocked in glycine (4 mg/mL). Larvae were then transferred into 4% formaldehyde at RT for 1 h. Hybridisation was carried out with DIG-labelled probes for 48 h at 60 °C. After incubation, samples were washed through serial dilutions of 25%, 50%, 75%, 100% 2x SSCT at hybridisation temperature. The colour development was carried out in a 1:50 dilution of NBT/BCIP at RT. Stained animals were visualised with a Leica DM1000 microscope equipped with a MC190 HD Microscope Camera (Leica, Germany). For each gene at least 30 specimens were tested. For double fluorescent ISH after the SSCT washes, samples were blocked in 0.5% blocking reagent (FP1020, PerkinElmer) at RT for 1 h. Then samples were incubated overnight with anti-digoxigenin (1:100) (Roche). After TNT (0.1M Tris-HCl pH 7.5/0.15M NaCl/0.5% Triton X-100) washes, samples were incubated in Cy3 (NEL744001KT, TSA Plus Kit, PerkinElmer). To stop the POD activity, the samples were washed in 0.1M glycine pH 2.0 and then incubated overnight with anti-fluorescein (1:250) (Roche). Samples were then washed in TNT, incubated with DAPI 1:1000. Samples were imaged on Leica TCS SP8 DLS confocal microscope.

### Whole-mount immunofluorescence

After fixation, the samples were washed 5 times with PBST (1× PBS, 0.05% (vol/vol) Tween-20) for 10 min. The samples were blocked in 5% BSA in PBST for 1 hr at RT. Primary antibody (1:500 dilution, mouse Anti-α-Tubulin Cat # T9026, Sigma-Aldrich) incubation was performed in a blocking solution (1% BSA in PBST) for 24-36 hr at 4°C. The samples were washed with PBST for 5 x 5 min, after which samples were incubated with secondary antibodies (1:250 dilution; Goat anti-Mouse IgG Alexa Fluor 594 Cat # A-11032, ThermoFisher) diluted in blocking solution for overnight at 4°C. Then, the samples were washed with PBST for 5 x 10 min. Imaging was performed on Leica TCS SP8 DLS and Leica DMi8 confocal microscopes.

### GPCR clustering and phylogeny

We used previously published GPCR datasets of *Crassostrea virginica, Platynereis dumerilii, Saccoglossus kovalevski* and *Drosophila melanogaster* [61], as well as *Homo sapiens* GPCRs [62]. Initially, to identify the *Nematostella* GPCR candidates grouping with bilaterians was done clustering analysis using CLANS [78] with the BLOSUM62 scoring matrix and a p-value cut-off of 1 x 10-25. Sequences from the main cluster were used in phylogenetic analysis. Sequences were aligned with MUSCLE [79] using default settings and trimmed with TrimAl using the automated mode [80]. Maximum likelihood phylogenetic analyses were completed using PhyML 3.0 online (http://www.atgc-montpellier.fr/phyml/) [81]. The model LG +G +F was automatically selected by the Smart Model Selection with (SH-aLRT) branch support with 1000 replicates.

## Supporting information

3D view of fluorescence in situ hybridisation for larval-specific neuron gene, related to Figure 5A. NVE8226 mRNA probe (yellow) and DAPI in Blue.

3D view of fluorescence in situ hybridisation for larval-specific neuron gene, related to Figure 5E. NVE14902 mRNA probe (yellow) and DAPI in Blue.

3D view of Double fluorescence in situ hybridisation for spot probe NVE14554 (green) and rind NVE8226 (red) mRNA probes, related to Figure 5J.

3D view of Double fluorescence in situ hybridisation for spot probe NVE14554 (green) and rind NVE14902 (red) mRNA probes, related to Figure 5M.

3D view of fluorescence in situ hybridisation for gland/secretory cells type 1, related to Figure 7B&C. gland/secretory cells type 1 marker NVE22589

3D view of fluorescence in situ hybridisation for gland/secretory cells type 3, related to Figure 7F. Gland/secretory cells type 3 marker NVE23674

Fig. S1: Additional neuronal associated genes DE between the apical and oral regions.

## Acknowledgments

We thank Kevin Atkins for his help with setting up the sea anemone facility. We also thank Alix Harvey and Yousef Touhami for their support in maintaining animal facility.

## Supplementary information

**Fig. S1:**
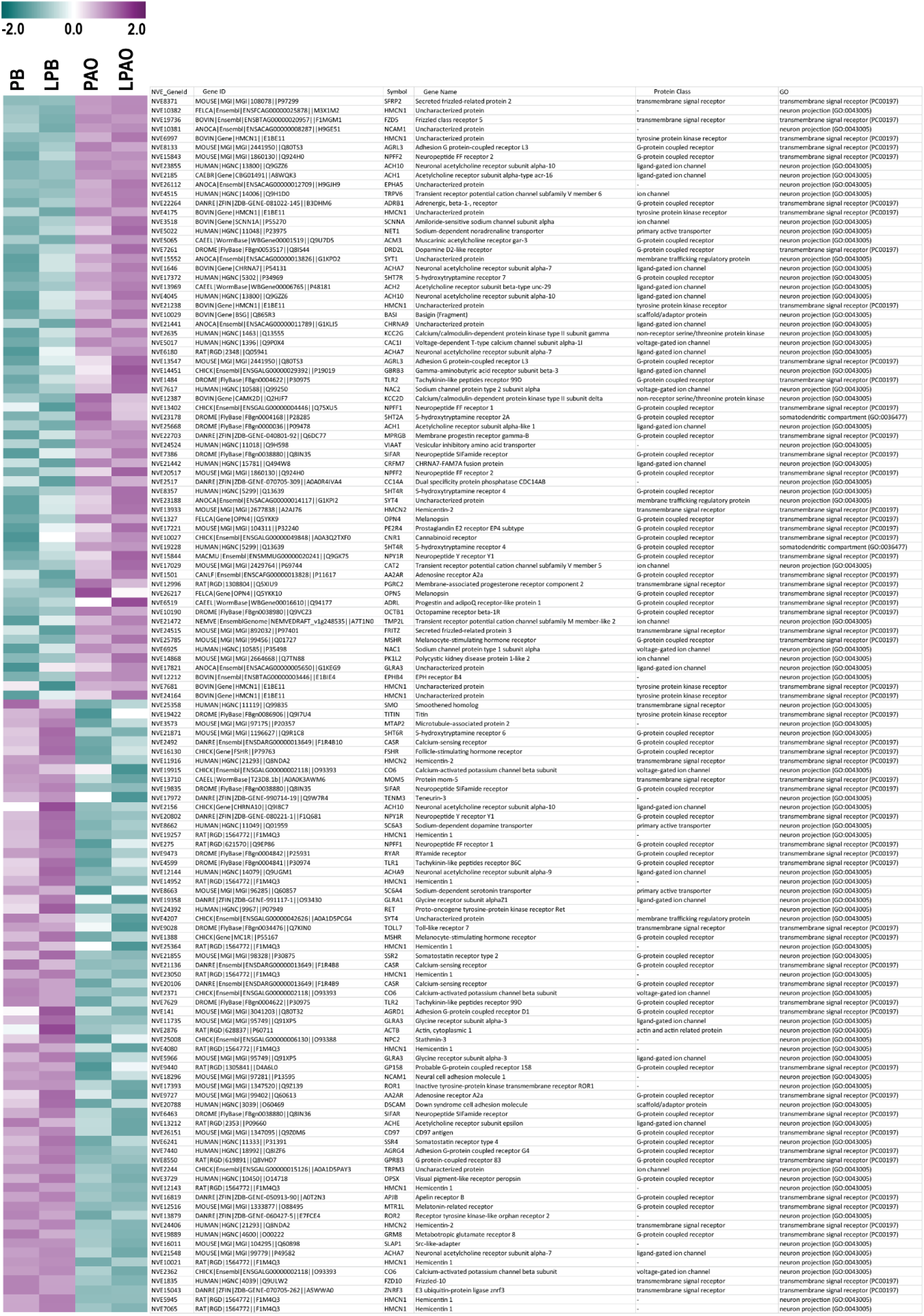
Additional neuronal associated genes DE between the apical and oral regions. Heatmap shows the relative expression of neuronal associated genes among oral or apical region of *Nematostella* larvae. The gene id and additional details are shown on the right.

**Table S1: Data relevant to figure2, 4, 6, 8 and 9**.

**Movie 1:** 3D view of fluorescence in situ hybridisation for larval-specific neuron gene, related to Figure 5A. *NVE8226* mRNA probe (yellow) and DAPI in Blue.

**Movie 2:** 3D view of fluorescence in situ hybridisation for larval-specific neuron gene, related to Figure 5E. *NVE14902* mRNA probe (yellow) and DAPI in Blue.

**Movie 3:** 3D view of Double fluorescence in situ hybridisation for spot probe *NVE14554* (green) and rind NVE8226 (red) mRNA probes, related to Figure 5J. DAPI in Blue.

**Movie 4:** 3D view of Double fluorescence in situ hybridisation for spot probe *NVE14554* (green) and rind *NVE14902* (red) mRNA probes, related to Figure 5M. DAPI in Blue.

**Movie 5:** 3D view of fluorescence in situ hybridisation for gland/secretory cells type 1, related to Figure 7B&C. gland/secretory cells type 1 marker NVE22589 (yellow) and DAPI in cyan.

**Movie 6:** 3D view of fluorescence in situ hybridisation for gland/secretory cells type 3, related to Figure 7F. Gland/secretory cells type 3 marker NVE23674 (yellow) and DAPI in cyan.

## References

1. Marinković, M., J. Berger, and G. Jékely, Neuronal coordination of motile cilia in locomotion and feeding. Philosophical Transactions of the Royal Society B: Biological Sciences, 2020. 375(1792): p. 20190165.

2. Sinigaglia, C., et al., Molecular characterization of the apical organ of the anthozoan Nematostella vectensis. Developmental biology, 2015. 398(1): p. 120–133.

3. Iwao, K., T. Fujisawa, and M. Hatta, A cnidarian neuropeptide of the GLWamide family induces metamorphosis of reef-building corals in the genus Acropora. Coral Reefs, 2002. c21: p. 127–129.

4. Schmich, J., S. Trepel, and T. Leitz, The role of GLWamides in metamorphosis of Hydractinia echinata. Dev Genes Evol, 1998. 208(5): p. 267–73.

5. Conzelmann, M., et al., Neuropeptides regulate swimming depth of *Platynereis* larvae. Proceedings of the National Academy of Sciences, 2011. 108(46): p. E1174–E1183.

6. Verasztó, C., et al., Ciliomotor circuitry underlying whole-body coordination of ciliary activity in the Platynereis larva. eLife, 2017. c6: p. e26000.

7. Goldberg, J.I., et al., Identification and evolutionary implications of neurotransmitter–ciliary interactions underlying the behavioral response to hypoxia in *Lymnaea stagnalis* embryos. The Journal of Experimental Biology, 2011. 214(16): p. 2660–2670.

8. Garner, S., et al., Neurogenesis in sea urchin embryos and the diversity of deuterostome neurogenic mechanisms. Development, 2016. c143: p. 286–297.

9. Gruhl, A., Serotonergic and FMRFamidergic nervous systems in gymnolaemate bryozoan larvae. Zoomorphology, 2009. c128: p. 135–156.

10. Page, L.R., Apical Sensory Organ in Larvae of the Patellogastropod Tectura scutum. Biological Bulletin, 2002. 202(1): p. 6–22.

11. Chia, F.-S. and R. Koss, Fine structural studies of the nervous system and the apical organ in the planula larva of the sea anemone Anthopleura elegantissima. Journal of Morphology, 1979. 160(3): p. 275–297.

12. Voronezhskaya, E.E., S.A. Tyurin, and L.P. Nezlin, Neuronal development in larval chiton Ischnochiton hakodadensis (Mollusca: Polyplacophora). Journal of Comparative Neurology, 2002. 444(1): p. 25–38.

13. Nezlin, L.P. and E.E. Voronezhskaya, Early peripheral sensory neurons in the development of trochozoan animals. Russian Journal of Developmental Biology, 2017. c48: p. 130–143.

14. Kumar, S., et al., The development of early pioneer neurons in the annelid Malacoceros fuliginosus. BMC Evolutionary Biology, 2020. 20(1): p. 117.

15. Marlow, H., et al., Larval body patterning and apical organs are conserved in animal evolution. BMC Biol, 2014. c12: p. 7.

16. Williams, E.A. and G. Jekely, Neuronal cell types in the annelid Platynereis dumerilii. Curr Opin Neurobiol, 2019. c56: p. 106–116.

17. Randel, N., et al., Expression dynamics and protein localization of rhabdomeric opsins in Platynereis larvae. Integr Comp Biol, 2013. 53(1): p. 7–16.

18. Veraszto, C., et al., Ciliary and rhabdomeric photoreceptor-cell circuits form a spectral depth gauge in marine zooplankton. Elife, 2018. 7.

19. Kelava, I., F. Rentzsch, and U. Technau, Evolution of eumetazoan nervous systems: insights from cnidarians. Philosophical Transactions of the Royal Society B: Biological Sciences, 2015. 370(1684): p. 20150065.

20. Marlow, H.Q., et al., Anatomy and development of the nervous system of Nematostella vectensis, an anthozoan cnidarian. Dev Neurobiol, 2009. 69(4): p. 235–54.

21. Layden, M.J., F. Rentzsch, and E. Röttinger, The rise of the starlet sea anemone Nematostella vectensis as a model system to investigate development and regeneration. WIREs Developmental Biology, 2016. 5(4): p. 408–428.

22. Zang, H. and N. Nakanishi, Expression Analysis of Cnidarian-Specific Neuropeptides in a Sea Anemone Unveils an Apical-Organ-Associated Nerve Net That Disintegrates at Metamorphosis. Front Endocrinol (Lausanne), 2020. c11: p. 63.

23. Piraino, S., et al., Complex neural architecture in the diploblastic larva of Clava multicornis (Hydrozoa, Cnidaria). J Comp Neurol, 2011. 519(10): p. 1931–51.

24. Gajewski, M., et al., LWamides from Cnidaria constitute a novel family of neuropeptides with morphogenetic activity. Roux’s archives of developmental biology, 1996. 205(5): p. 232–242.

25. Leitz, T. and M. Lay, Metamorphosin A is a neuropeptide. Roux’s archives of developmental biology, 1995. 204(4): p. 276–279.

26. Katsukura, Y., et al., Control of planula migration by LWamide and RFamide neuropeptides in *Hydractinia echinata*. Journal of Experimental Biology, 2004. 207(11): p. 1803–1810.

27. Woollacott, R.M., Structure and swimming behavior of the larva of Haliclona tubifera (Porifera: Demospongiae). Journal of Morphology, 1993. 218(3): p. 301–321.

28. Richards, G.S. and B.M. Degnan, The expression of Delta ligands in the sponge Amphimedon queenslandica suggests an ancient role for Notch signaling in metazoan development. EvoDevo, 2012. 3(1): p. 15.

29. Leys, S.P. and B.M. Degnan, Cytological Basis of Photoresponsive Behavior in a Sponge Larva. The Biological Bulletin, 2001. 201(3): p. 323–338.

30. Maldonado, M., et al., The cellular basis of photobehavior in the tufted parenchymella larva of demosponges. Marine Biology, 2003. 143(3): p. 427–441.

31. Jekely, G., Origin and early evolution of neural circuits for the control of ciliary locomotion. Proc Biol Sci, 2011. 278(1707): p. 914–22.

32. Degnan, S.M. and B.M. Degnan, The origin of the pelagobenthic metazoan life cycle: what’s sex got to do with it? Integrative and Comparative Biology, 2006. 46(6): p. 683–690.

33. Nakanishi, N., et al., Sensory Flask Cells in Sponge Larvae Regulate Metamorphosis via Calcium Signaling. Integrative and Comparative Biology, 2015. 55(6): p. 1018–1027.

34. Ueda, N., et al., An ancient role for nitric oxide in regulating the animal pelagobenthic life cycle: evidence from a marine sponge. Scientific Reports, 2016. 6(1): p. 37546.

35. Nielsen, C., Life cycle evolution: was the eumetazoan ancestor a holopelagic, planktotrophic gastraea? BMC Evol Biol, 2013. c13: p. 171.

36. Nielsen, C., Larval and adult brains1. Evolution & Development, 2005. 7(5): p. 483–489.

37. Sinigaglia, C., et al., The bilaterian head patterning gene six3/6 controls aboral domain development in a cnidarian. PLoS Biol, 2013. 11(2): p. e1001488.

38. Matus, D.Q., et al., Molecular evidence for deep evolutionary roots of bilaterality in animal development. Proc Natl Acad Sci U S A, 2006. 103(30): p. 11195–200.

39. Arendt, D., M.A. Tosches, and H. Marlow, From nerve net to nerve ring, nerve cord and brain--evolution of the nervous system. Nat Rev Neurosci, 2016. 17(1): p. 61–72.

40. Sebe-Pedros, A., et al., Cnidarian Cell Type Diversity and Regulation Revealed by Whole-Organism Single-Cell RNA-Seq. Cell, 2018. 173(6): p. 1520–1534 e20.

41. Sigg, M.A., et al., Evolutionary Proteomics Uncovers Ancient Associations of Cilia with Signaling Pathways. Developmental Cell, 2017. 43(6): p. 744-762.e11.

42. Pennati, R., et al., Neural system reorganization during metamorphosis in the planula larva of Clava multicornis (Hydrozoa, Cnidaria). Zoomorphology, 2013. 132(3): p. 227–237.

43. Nakanishi, N., et al., Early development, pattern, and reorganization of the planula nervous system in Aurelia (Cnidaria, Scyphozoa). Dev Genes Evol, 2008. 218(10): p. 511–24.

44. Martin, V.J. and F.-S. Chia, Fine Structure of a Scyphozoan Planula, Cassiopeia xamachana. Biological Bulletin, 1982. 163(2): p. 320–328.

45. Steinmetz, P.R.H., et al., Gut-like ectodermal tissue in a sea anemone challenges germ layer homology. Nat Ecol Evol, 2017. 1(10): p. 1535–1542.

46. Babonis, L.S., et al., Genomic analysis of the tryptome reveals molecular mechanisms of gland cell evolution. Evodevo, 2019. c10: p. 23.

47. Sachkova, M.Y., et al., The Birth and Death of Toxins with Distinct Functions: A Case Study in the Sea Anemone Nematostella. Mol Biol Evol, 2019. 36(9): p. 2001–2012.

48. Columbus-Shenkar, Y.Y., et al., Dynamics of venom composition across a complex life cycle. Elife, 2018. 7.

49. Leys, S.P., et al., Spectral sensitivity in a sponge larva. J Comp Physiol A Neuroethol Sens Neural Behav Physiol, 2002. 188(3): p. 199–202.

50. Collin, R., et al., Phototactic responses of larvae from the marine sponges Neopetrosia proxima and Xestospongia bocatorensis (Haplosclerida: Petrosiidae). Invertebrate Biology, 2010. 129(2): p. 121–128.

51. Soliman, S., Pharmacological control of ciliary activity in the young sea urchin larva. Effects of monoaminergic agents. Comparative Biochemistry and Physiology Part C: Comparative Pharmacology, 1983. 76(1): p. 181–191.

52. Soliman, S., Pharmacological control of ciliary activity in the young sea urchin larva. Studies on the role of Ca2+ and cyclic nucleotides. Comparative Biochemistry and Physiology Part C: Comparative Pharmacology, 1984. 78(1): p. 183–191.

53. Kuang, S., et al., Serotonergic sensory-motor neurons mediate a behavioral response to hypoxia in pond snail embryos. Journal of Neurobiology, 2002. 52(1): p. 73–83.

54. Lacalli, T.C., T.H.J. Gilmour, and J.E. West, Ciliary band innervation in the bipinnaria larva of <i>Pisaster ochraceus</i>. Philosophical Transactions of the Royal Society of London. Series B: Biological Sciences, 1990. 330(1258): p. 371–390.

55. Lacalli, T.C. and T.H.J. Gilmour, Ciliary reversal and locomotory control in the pluteus larva of <i>Lytechinus pictus</i>. Philosophical Transactions of the Royal Society of London. Series B: Biological Sciences, 1990. 330(1258): p. 391–396.

56. Wada, Y., Y. Mogami, and S. Baba, Modification of ciliary beating in sea urchin larvae induced by neurotransmitters: beat-plane rotation and control of frequency fluctuation. J Exp Biol, 1997. 200(Pt 1): p. 9–18.

57. Bergles, D. and S. Tamm, Control of Cilia in the Branchial Basket of Ciona intestinalis (Ascidacea). The Biological Bulletin, 1992. 182(3): p. 382–390.

58. Alvarez, L., et al., The rate of change in Ca2+ concentration controls sperm chemotaxis. Journal of Cell Biology, 2012. 196(5): p. 653–663.

59. Tamm, S.L. and S. Tamm, Ciliary reversal without rotation of axonemal structures in ctenophore comb plates. Journal of Cell Biology, 1981. 89(3): p. 495–509.

60. Hewes, R.S. and P.H. Taghert, Neuropeptides and neuropeptide receptors in the Drosophila melanogaster genome. Genome Res, 2001. 11(6): p. 1126–42.

61. Thiel, D., et al., Nemertean, brachiopod and phoronid neuropeptidomics reveals ancestral spiralian signalling systems. bioRxiv, 2021: p. 2021.03.03.433790.

62. Quiroga Artigas, G., et al., A G protein–coupled receptor mediates neuropeptide-induced oocyte maturation in the jellyfish Clytia. PLOS Biology, 2020. 18(3): p. e3000614.

63. Yaguchi, S., et al., ankAT-1 is a novel gene mediating the apical tuft formation in the sea urchin embryo. Developmental Biology, 2010. 348(1): p. 67–75.

64. Dictus, W.J.A.G. and P. Damen, Cell-lineage and clonal-contribution map of the trochophore larva of Patella vulgata (Mollusca)1Both authors contributed equally to this work.1. Mechanisms of Development, 1997. 62(2): p. 213–226.

65. Arenas-Mena, C., K.S.-Y. Wong, and N. Arandi-Forosani, Ciliary band gene expression patterns in the embryo and trochophore larva of an indirectly developing polychaete. Gene Expression Patterns, 2007. 7(5): p. 544–549.

66. Bloodgood, R.A., Sensory reception is an attribute of both primary cilia and motile cilia. Journal of Cell Science, 2010. 123(4): p. 505–509.

67. Shah, A.S., et al., Motile Cilia of Human Airway Epithelia Are Chemosensory. Science, 2009. 325(5944): p. 1131–1134.

68. Genikhovich, G. and U. Technau, Induction of spawning in the starlet sea anemone Nematostella vectensis, in vitro fertilization of gametes, and dejellying of zygotes. Cold Spring Harb Protoc, 2009. 2009(9): p. ppdb prot5281.

69. Bolger, A.M., M. Lohse, and B. Usadel, Trimmomatic: a flexible trimmer for Illumina sequence data. Bioinformatics, 2014. 30(15): p. 2114–2120.

70. S, A., FastQC: a quality control tool for high throughput sequence data. Available online at: http://www.bioinformatics.babraham.ac.uk/projects/fastqc. 2010.

71. Langmead, B. and S.L. Salzberg, Fast gapped-read alignment with Bowtie 2. Nature methods, 2012. 9(4): p. 357–359.

72. Love, M.I., W. Huber, and S. Anders, Moderated estimation of fold change and dispersion for RNA-seq data with DESeq2. Genome Biology, 2014. 15(12): p. 550.

73. Altschul, S.F., et al., Gapped BLAST and PSI-BLAST: a new generation of protein database search programs. Nucleic Acids Research, 1997. 25(17): p. 3389–3402.

74. Mi, H., A. Muruganujan, and P.D. Thomas, PANTHER in 2013: modeling the evolution of gene function, and other gene attributes, in the context of phylogenetic trees. Nucleic Acids Research, 2012. 41(D1): p. D377–D386.

75. Huang da, W., B.T. Sherman, and R.A. Lempicki, Bioinformatics enrichment tools: paths toward the comprehensive functional analysis of large gene lists. Nucleic Acids Res, 2009. 37(1): p. 1–13.

76. Wolenski, F.S., et al., Characterizing the spatiotemporal expression of RNAs and proteins in the starlet sea anemone, Nematostella vectensis. Nature Protocols, 2013. 8(5): p. 900–915.

77. Genikhovich, G. and U. Technau, In Situ Hybridization of Starlet Sea Anemone (Nematostella vectensis) Embryos, Larvae, and Polyps. Cold Spring Harbor Protocols, 2009. 2009(9): p. ppdb.prot5282.

78. Frickey, T. and A. Lupas, CLANS: a Java application for visualizing protein families based on pairwise similarity. Bioinformatics, 2004. 20(18): p. 3702–4.

79. Edgar, R.C., MUSCLE: a multiple sequence alignment method with reduced time and space complexity. BMC Bioinformatics, 2004. c5: p. 113.

80. Capella-Gutierrez, S., J.M. Silla-Martinez, and T. Gabaldon, trimAl: a tool for automated alignment trimming in large-scale phylogenetic analyses. Bioinformatics, 2009. 25(15): p. 1972–3.

81. Guindon, S., et al., New algorithms and methods to estimate maximum-likelihood phylogenies: assessing the performance of PhyML 3.0. Syst Biol, 2010. 59(3): p. 307–21.

